# RO8191, a new compound for initiating embryo implantation in mice

**DOI:** 10.1101/2024.09.30.615945

**Authors:** Junlan Shu, Jumpei Terakawa, Satoko Osuka, Ayako Muraoka, Jiali Ruan, Junya Ito, Atsuo Iida, Eiichi Hondo

## Abstract

During early pregnancy in mice, leukemia inhibitory factor (LIF) regulates embryo implantation by activating the JAK/STAT3 signaling pathway. The STAT3 pathway has been recognized to play a critical role in embryo implantation. However, it is not clear whether STAT3 activation itself can cause induction of embryo implantation. In this study, the effects of RO8191, a potential STAT3 activator, on embryo implantation were investigated through a series of studies with different mouse models. We found that the injection of RO8191 can induce embryo implantation by activating the STAT3 pathway in delayed implantation mice. Furthermore, RO8191 can initiate decidualization, which is essential for embryo implantation, in uterine epithelial-specific *Stat3, Gp130*, or *Lifr* conditional knockout (cKO) mice, which shows infertility due to embryo implantation failure. Histomorphological observations revealed that successful embryo implantation and embryonic development was seen in *Lifr* cKO mice. Increased epithelial detachment and vascularization were observed in *Stat3* cKO mice, and excessive inflammatory response and embryo death were observed in *Gp130* cKO mice. These results suggest that STAT3, Gp130 and LIFR each play a distinct role in embryo implantation and development. Although the specific mechanisms of RO8191 are not fully understood, the present study will provide insights to support the application of RO8191 in treating recuurent implantation failure.

## Introduction

Embryo implantation is a crucial stage in early mammalian gestation. The blastocyst hatches, attaches, adheres, and invades the receptive uterus in humans and mice (Wang and Dey 2006). Failure of embryo implantation is a significant challenge in the field of infertility treatment. Previous studies have shown that the success rate of implantation per embryo hovers around 25% in human populations with normal fertility, according to both natural cycles and assisted reproductive technology indications (Macklon, Geraedts, and Fauser 2002). Although the exact molecular mechanisms of blastocyst implantation are not fully understood, existing research has revealed that successful implantation depends on the synchronization of endometrial receptivity with blastocyst activation (Giudice 1999). For example, insufficient endometrial receptivity can lead to recurrent implantation failure (RIF) in clinical pregnancies (Coughlan *et al*. 2014; Kliman and Frankfurter 2019). Endometrial receptivity is controlled by the hormones 17β-estradiol (E2) and progesterone (P4) (Wang and Dey 2006), which are primarily synthesized by the ovaries.

In mice, embryo implantation occurs on the fourth day of pregnancy (D4) (D1= the day the vaginal plug is found). This process is triggered by the transient elevation of E2 (Kojima, Tam, and Tam 2014; Yang *et al*. 1995). E2 can stimulate the release of leukemia inhibitory factor (LIF), a cytokine that belongs to the interleukin (IL)-6 family cytokines (including oncostatin M, IL-11, IL-27, ciliary neurotrophic factor [CNTF], and cardiotrophin-1 [CT-1]) (Hu *et al*. 2007). It is widely accepted that LIF is expressed in the uterine glandular epithelium (GE) during the pre-implantation phase (Song *et al*. 2000) and is considered crucial for implantation (Robb *et al*. 2002). The LIF interacts with the LIF receptor (LIFR) on the luminal epithelium and forms a heterodimer with the glycoprotein 130 (Gp130) (Onishi and Zandstra 2015). This complex heterodimer activates the JAK/STAT3 signaling pathway, leading to the phosphorylation of Janus kinase and subsequent phosphorylation of STAT3 (Pastuschek *et al*. 2015; Salleh 2014). Following phosphorylation, p-STAT3 translocates to the nucleus, where it mediates the expression of a variety of genes in the cell (Onishi and Zandstra 2015).

Infertility due to failure of embryo adhesion has been observed in *Lif*-deficient mice (Stewart *et al*. 1992). The importance of LIF for embryo implantation has been further demonstrated by the finding that intraperitoneal administration of a LIF antagonist (White et al. 2007) or anti-LIF antibody (Terakawa *et al*. 2011) in the C57BL/6J (B6) mouse strain prevented embryo implantation, leading to infertility. Intriguingly, injection of recombinant LIF or recombinant CT-1 can induce embryo implantation by activating STAT3 signaling in the uterine epithelium in delayed implantation (DI) mice (ICR or B6 strain) (Kobayashi *et al*. 2014). In contrast, CNTF, which belongs to the same IL-6 cytokine family as LIF and CT-1, neither induced embryo implantation nor activated STAT3 phosphorylation in DI mice. Mice with a conditional knockout of *Stat3* in the uterine epithelium exhibit infertility due to implantation failure (Hiraoka *et al*. 2020). Similarly, mice with a conditional knockout of either *Lifr* (Fukui *et al*. 2021) or *Gp130* (Namiki *et al*. 2023) in the uterine epithelium are also infertile due to failure of blastocyst implantation. These findings suggest that the epithelial-originated LIFR/Gp130-JAK/STAT3 signaling pathway is essential for successful embryo implantation in mice.

Successful embryo implantation can be achieved through activation of STAT3, but to date, no pharmacological agents have been used to activate the STAT3 pathway and induce implantation in mice. The purpose of the present study was to determine whether the pharmacological agents could induce embryo implantation by activating the JAK/STAT3 signaling pathway. We found that RO8191, an interferon-α receptor 2 agonist (Konishi et al. 2012), is a useful STAT3 activator that can induce embryo implantation.

## Materials and methods

### Animals

This study was approved by the Ethics Review Board for Animal Experiments at Nagoya University (approval number: A240034-002) and the Ethical Committee for Vertebrate Experiments at Azabu University (approval number: 230327-9). All experiments were conducted in accord with the relevant guidelines and regulations, including the Animal Research: Reporting of In Vivo Experiments (ARRIVE) guidelines.

ICR (Japan SLC, Shizuoka, Japan), C57BL/6J (Jackson Laboratory Japan, Kanagawa, Japan), and the following mouse strains, all aged over seven weeks, were used in the experiments: *Ltf*^*iCre/+*^ mouse < *Ltf*^*tm1(icre)Tdku*^*/J*, JAX: 026030 > (Daikoku et al. 2014), *Stat3*^*flox/flox*^ (*Stat3*^*f/f*^) <*Stat3*^*tm2Aki*^> (Takeda et al. 1998), *Gp130*^*flox/flox*^ (*Gp130*^*f/f*^) mouse <*Il6st*^*tm1Wme*^> (Betz et al. 1998), and *Lifr*^*flox/flox*^ (*Lifr*^*f/f*^) <*Lifr*^*tm1c(EUCOMM)Hmgu*^>. To obtain the *Lifr*^*flox/flox*^ mice (tm1c), *Lifr*^*tm1a(EUCOMM)Hmgu*^ purchased from the European Mouse Mutant Archive (EMMA) (strain ID: EM:06941), harboring a knockout first allele, were crossed with FLPe transgenic mice <Tg(CAG-flpe)36Ito> (RBRC01834) (Kanki, Suzuki, and Itohara 2006) and FRT-LacZ-neo-FRT cassette was removed and loxP-flanked exon 4 was left. *Stat3*^*f/f*^, *Gp130*^*f/f*^ and *Lifr*^*f/f*^ mice were crossed with *Ltf*^*iCre/+*^ mouse to generate uterine epithelial-specific gene deficient (conditional knockout, cKO) (*Stat3* cKO, *Gp130* cKO or *Lifr* cKO) mice as previously described (Namiki et al. 2023), respectively. The primers used for genotyping were; 5′-GTTTCCTCCTTCTGGGCTCC-3′, 5′-TTTAGTGCCCAGCTTCCCAG-3′ and 5′-CCTGTTGTTCAGCTTGCACC-3′ for *Ltf*^*iCre*^; 5′-CCTGAAGACCAAGTTCATCTGTGTGAC-3′, 5′-CACACAAGCCATCAAACTCTGGTCTCC-3′ and 5′-GATTTGAGTCAGGGATCCTTATCTTCG-3′ for *Stat3*^*flox*^; 5′-GGCTTTTCCTCTGGTTCTTG-3′ and 5′-CAGGAACATTAGGCCAGATG-3′ for *Gp130*^*flox*^; 5′-TGAGAGCACGGAAGCTCTTT-3′ and 5′-ACTGCCCGACAAGGTTTTTA-3′ for *Lifr*^*flox*^.

All the mouse strains, except ICR, were maintained in a C57BL/6 strain background and housed in the barrier facility at Azabu University. ICR mice were housed in the barrier facility at Nagoya University. The mice were fed *ad libitum* under controlled light-dark cycles (12 hours of light followed by 12 hours of dark) at 22 ± 3 °C. The first day of pregnancy (D1) was determined as the morning when a vaginal plug was observed in the female that had been mated with fertile wildtype males on the previous evening.

### RO8191 treatment in DI mice

The experimental procedure is shown in Figure 1A. To induce delayed implantation (DI) model mice, plug-positive ICR females were ovariectomized between 1300 and 1530 h on D3 under sevoflurane (193-17791, FujiFilm Wako Pure Chemicals, Osaka, Japan) anesthesia, and medroxyprogesterone acetate (100 μl/head; Pfizer Inc, NY, USA) was injected subcutaneously in the ventral region. The RO8191 (T22142, TargetMol) was dissolved in sesame oil (196-15385, FujiFilm Wako Pure Chemicals), and a single intraperitoneal (i.p.) injection of RO8191 (400 μg/head; dissolved in sesame oil) or sesame oil was performed at 1300 h on D7. These female mice were euthanized in a carbon dioxide chamber, followed by dissection at 1300 h on D10 to count the number of implantation sites. The saline solution was used to flush the uterine horns to check for the presence of embryos if the uteri did not show the implantation sites. DI mice were excluded from the statistical analysis if they had no implantation sites and no blastocyst was recovered. ICR females injected with sesame oil were used as a control. The implantation rates of two groups (oil and RO8191), which were defined as the ratio of the number of mice with implantation sites to the total number of mice after being injected with oil or RO8191.

**Fig. 1.**
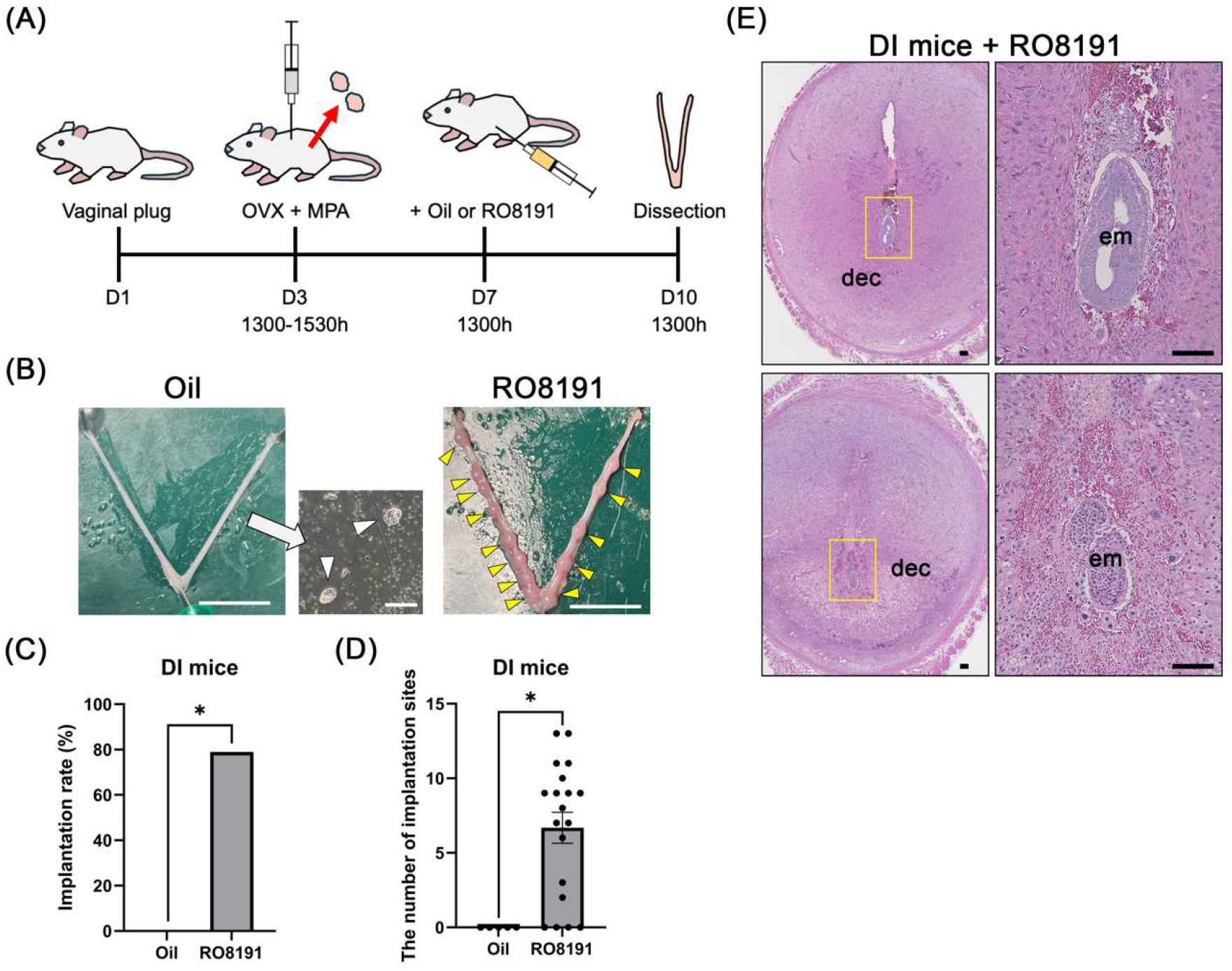
RO8191 induces embryo implantation in delayed implantation (DI) model mice. **(A)** The experimental procedure of the artificial DI mouse. Ovariectomy (OVX) and subcutaneous administration of siliconized medroxyprogesterone acetate (MPA) were performed on D3. A single injection of oil or RO8191 was performed on D7. Females were euthanized and dissected on D10. **(B)** Representative images of the gross uterine morphology. Blastocysts were recovered from oil-treated mice (white arrowheads). Implantation sites (yellow arrowheads on right panel) were visible in DI mice after RO8191 treatment. Scale bars: 1 cm for uterine images and 200 μm for blastocysts. **(C, D)** Implantation rate (C) and the number of implantation sites (D) in the oil and RO8191-treated DI mice. * indicates significantly different (p < 0.0001). **(E)** Representative histological image of uterine cross-sections of implantation sites on D10 after RO8191 treatment in DI mice. Right panels are higher magnification of left panels indicated by yellow line. Lower panels indicated abnormal embryo development and decidual reaction. Scale bars: 100 μm. dec: decidua; em: embryo.

### RO8191 treatment in conditional knockout mice

The experimental procedure is shown in Figure 3A. Plug-positive females of *Stat3, Gp130* or *Lifr* cKO were administered with a single i.p. injection of RO8191 (400 μg/head; dissolved in sesame oil) or sesame oil at between 1330 and 1700 h on D4. Plug-positive females of *Stat3*^*f/f*^, *Gp130*^*f/f*^ or *Lifr* ^*f/f*^ were injected with sesame oil as a pregnant control group. The number of implantation sites was counted macroscopically on D7 or D8. If there was no implantation site, the uterine horns were flushed with saline solution to retrieve the embryos and determine the success of mating. Mice were excluded from the statistical analysis, if they had no implantation sites and no blastocyst was recovered. Images were saved in PNG format at a resolution of 806×1441 pixels. ImageJ software (version 1.54f; National Institutes of Health, Bethesda, MD, USA) was utilized to measure the areas of uterine bulges. The scale was set using the ‘Set Scale’ function based on a 1 cm scale bar. The bulges were selected using ‘Oval selection’. The measurements were recorded in square centimeters (cm^2^). The size of each bulge was measured in the control and cKOs (*Stat3, Gp130*, and *Lifr*) uteri on D7. Statistical analysis was conducted using GraphPad Prism (version 10.0; GraphPad Software), with results expressed as mean ± standard error of the mean (SEM). One-way ANOVA followed by Tukey’s multiple comparison test was used to assess group differences, with significance set at p < 0.05. In this study, we defined the bulges in the cKOs uterus as the putative implantation site.

### Isolation of LE

After the injection of RO8191 or sesame oil for 24 hours, DI mice were sacrificed by inhalation of carbon dioxide, and the uterine horns were collected. For the sesame oil injection group, each uterine horn was rinsed with 0.5 mL saline solution to recover blastocysts, while the uteri without embryos were excluded from sampling and discarded. The uteri were washed three times briefly with saline solution on ice. All uterine horns were opened longitudinally and cut into 2 mm pieces. The uterine pieces were collected in a tube and kept at −80°C overnight and incubated with RNA stabilization solution (AM7020, Thermo Fisher Scientific, Waltham, MA, USA) at −80°C for 1 hour. The luminal epithelium (LE) was isolated from the uterine pieces using forceps on the inner surface of the endometrium and collected by gravity sedimentation, washed in RNA stabilization solution, and stored at −80°C prior to protein extraction.

### Protein extraction and Western blot analysis

The RIPA lysis buffer containing a 1% protease inhibitor cocktail (25955-24; Nacalai tesque, Kyoto, Japan) was used for total protein isolation. Liver tissue was obtained from nonpregnant ICR mice. The LE or liver was lysed for 5 minutes on ice and centrifuged at 15,000 rpm for 30 minutes at 4°C for protein extraction. Total protein concentrations were determined using the Qubit™ protein assay kit (2411539, Thermo Fisher Scientific). LE protein (30 μg) and liver protein (10 μg) were separated on an 8% sodium dodecyl sulfate-polyacrylamide gel (SDS-PAGE) and transferred onto nitrocellulose membranes, which were followed by the blocking with 5% nonfat dry milk (Megmilk Snow Brand, Tokyo, Japan)/TTBS (Tris buffered saline with 0.01%-detergent) for 30 minutes at room temperature (RT). The membranes were then incubated with a primary antibody dilution buffer, consisting of TTBS diluted with 5% bovine serum albumin (010-25783, FujiFilm Wako Pure Chemicals) and primary antibodies, either STAT3 (1:2,000, 79D7, Cell Signaling Technology, Danvers, MA, USA) or phospho-STAT3 (p-STAT3 (Tyr705), 1:2,000, Cell Signaling Technology). After six 5-minte washes in TTBS solution, the membranes were incubated with horseradish peroxidase (HRP) conjugated goat anti-rabbit antibody (PI-1000-1, Vector Labs, Newark, CA, USA) for 1 hour at RT, followed by six 5-minte washes in. The Luminata™ Crescendo Western HRP Substrate (WBLUR0500, Merck Millipore, Burlington, MA, USA) immersed nitrocellulose membrane was placed in the Lumicube (Liponics, Tokyo, Japan), the aperture was adjusted to f/2.8, and the ISO was set to 12,800 with an exposure time of 30 seconds for photography.

### Histological methods

Uterine tissues from the mice were fixed in 4% paraformaldehyde (PFA) solution in phosphate-buffered saline (PBS) (pH 7.4) using standard histological techniques (Presnell and Schreibman, 2013; Suvarna et al., 2013). These tissues were dehydrated through increasing concentrations of ethanol solutions, cleared with xylene, and embedded in paraffin blocks. A plane of transverse sections was cut at 5 μm thickness and stained with the routine hematoxylin and eosin (H&E). Micrographs were captured by BZ-X700 microscopy (Keyence, Osaka, Japan).

### Statistical analysis

The parameter data including the implantation sites and implantation rate in the DI and cKOs mice were expressed as mean ± SEM or mean. Statistical analysis was performed by using independent Student’s t test at the GraphPad Prism (version 10.0; GraphPad Software). P values less than 0.05 were considered statistically significant.

## Results

### RO8191 promotes blastocysts implantation in DI mice

RO8191 has been reported to bind to the interferon-α receptor 2 and promote phosphorylation of STAT3 (Konishi et al. 2012). To validate the promotion activity of RO8191 for implantation in mice, RO8191 was injected into the DI mice (Fig. 1A). No implantation site was detected in the control mice, and nonadherent blastocysts were recovered after washing the uterus lumen (n = 5) (Fig. 1B). On the other hand, distinct implantation sites were observed in approximately 80% (15/19) of the RO8191-injected DI mice (Fig. 1B-D). In the negative samples after RO8191 injection (4/19), no implantation site was found, and the blastocysts were recovered. We confirmed the growing embryo in the implantation sites in RO8191-treated mice by the histological observation (Fig. 1E, upper panels), although some part of implantation sites showed abnormal embryo development and decidual reaction (Fig. 1E, lower panels). The implantation rate was significantly different between the two groups (Fig. 1C). The number of average implantation sites was 6.7 ± 1.0 in RO8191 group (Fig. 1D).

### Phosphorylation of STAT3 by RO8191

To evaluate whether RO8191 initiates implantation by regulating the JAK/STAT3 signaling pathway, the LE cells was collected from DI mice 24 hours after RO8191 injection. STAT3 expression was comparable between the control and RO8191-injected mice by Western blot analysis (Fig. 2). Phosphorylated-STAT3 (p-STAT3) was detectable after the intraperitoneal injection of RO8191 in DI mice compared to the control. High expression levels of both STAT3 and p-STAT3 were observed in the liver tissue from nonpregnant ICR mice as a control. LE cells only expressed the STAT3α isoform (86 kDa), whereas both STAT3α (86 kDa) and STAT3β (79 kDa) isoforms were expressed in the liver.

**Fig. 2.**
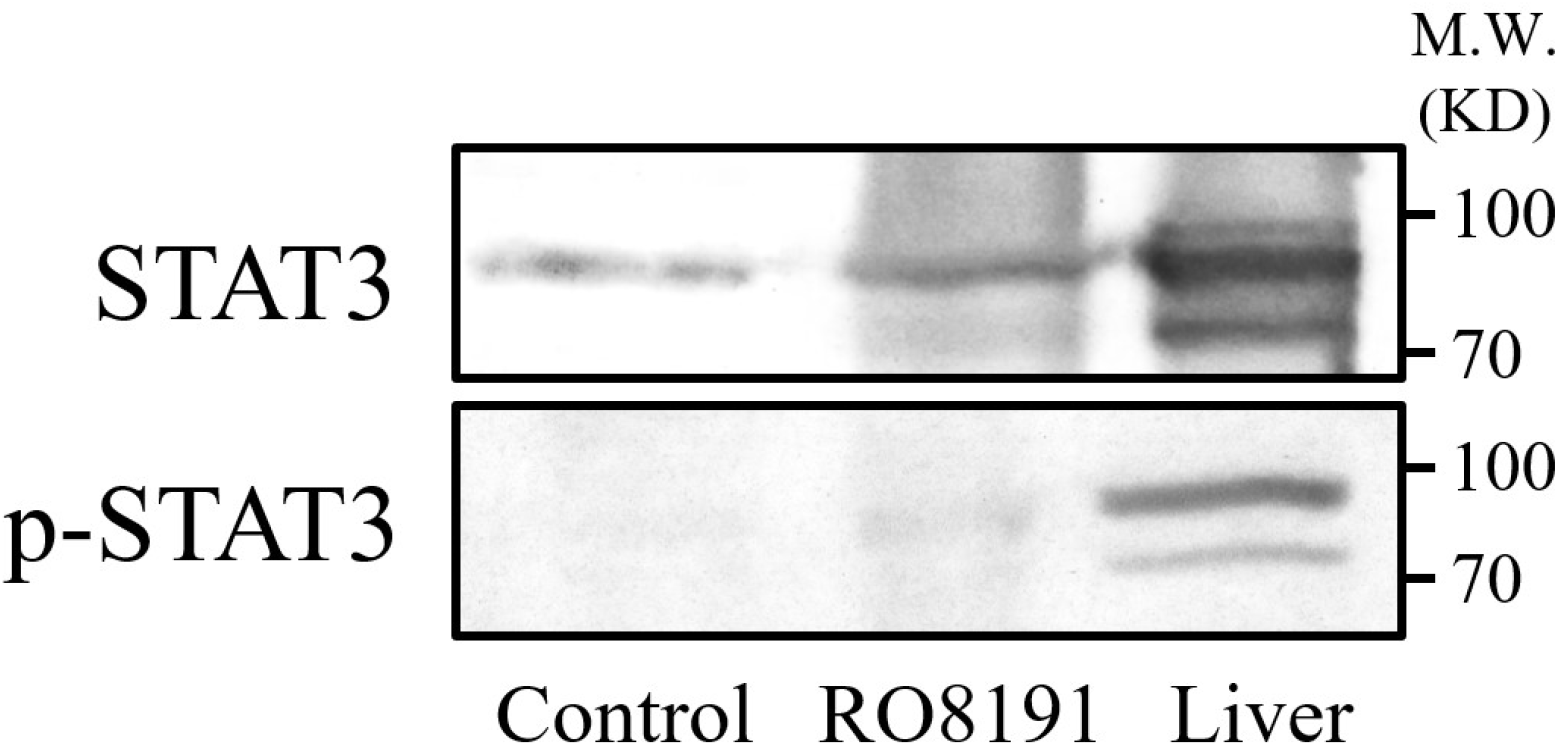
Western blot analysis of STAT3 and p-STAT3 for the LE cells after RO8191 treatment. Band is present at 86 and 79 kDa.

**Fig. 3.**
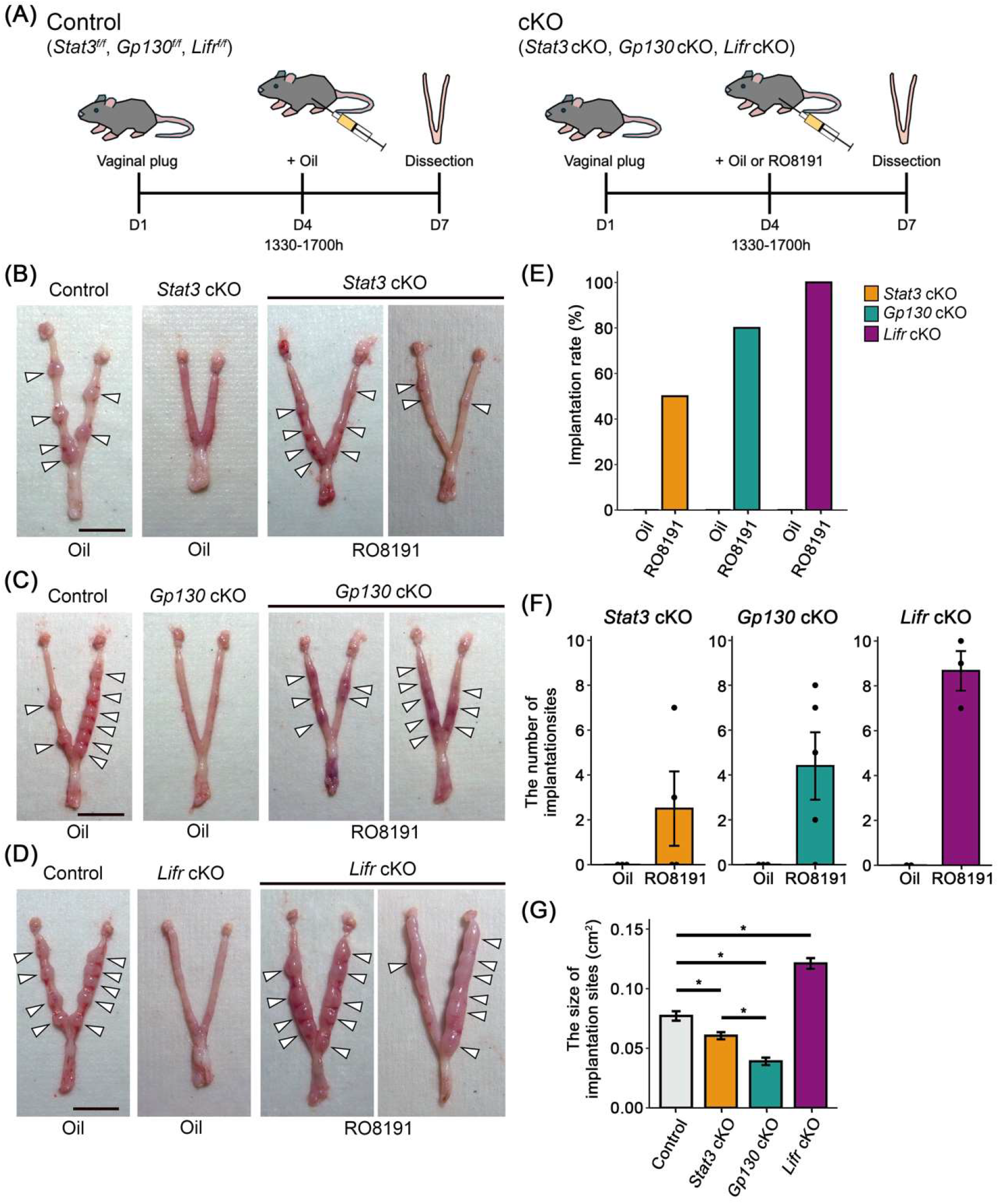
RO8191 induces decidualization in mice with genetic implantation failure. **(A)** The experimental procedure of the genetically modified mice. For control, a single oil injection was performed on D4. For cKOs, a single injection of oil or RO8191 was performed on D4. Females were euthanized and dissected on D7. (B-D) Uterine morphology of cKOs mice after the injection of oil or RO8191. The white arrowheads indicate the presumptive implantation site. Scale bars: 1 cm. **(E)** The implantation rate in cKOs mice after oil or RO8191 administration. **(F)** The number of implantation sites in cKOs mice after the injection of oil or RO8191. **(G)** The size of implantation sites in cKOs mice after oil or RO8191 administration. * indicates significantly different (p < 0.05).

### The effects on RO8191 for cKOs mice

Since the LIF-LIFR/Gp130-JAK/STAT3 signaling pathway is critical for establish for establishing embryo implantation in mice, RO8191 was administered to *Stat3* cKO (Hiraoka et al. 2020), *Gp130* cKO (Namiki et al. 2023) and *Lifr* cKO (Cheng et al. 2017; Fukui et al. 2021) mice to examine whether their implantation failure phenotype can be restored (Fig. 3A). As a pregnant control, *Stat3*^*f/f*^, *Gp130*^*f/f*^ and *Lifr*^*f/f*^ mice were injected with the same dose of sesame oil (Fig. 3A and Table 1).

**Table 1.** The percentage of implantation rate and the number of implantation sites of the flox mice.

| Strain | Injection | Implantation/n | The number of implantation sites<br>(mean ± SEM) |
| --- | --- | --- | --- |
| *Stat3<sup>ff</sup>* | Oil | 5/5 | 5.6 ± 0.8 |
| *Gp130<sup>ff</sup>* | | 3/3 | 9.7 ± 0.9 |
| *Lifr<sup>ff</sup>* | | 4/4 | 8.3 ± 0.5 |

In the *Stat3*^*f/f*^ (n = 5), *Gp130*^*f/f*^ (n = 3), and *Lifr*^*f/f*^ (n = 4) mice after oil injection on D4, implantation sites were lined along with the uteri on D7 and no defect of embryo implantation was found in each mouse strain (Fig. 3A-D and Table 1). The average number of implantation sites was 5.6, 9.7, and 8.3 in *Stat3*^*f/f*^, *Gp130*^*f/f*^ and *Lifr*^*f/f*^ mice, respectively (Table 1). Implantation sites, including small uterine bulges, were observed in *Stat3* cKO (2/4), *Gp130* cKO (4/5), and *Lifr* cKO (3/3) cKO mice injected with RO8191 (Fig. 3B-F). The average number of implantation sites was 2.5, 4.4, and 8.7 in *Stat3, Gp130*, and *Lifr* cKO mice, respectively (Fig. 3F). Administration of sesame oil in cKOs mice did not promote implantation (Fig. 3B-F). The size of the implantation bulges varied among the different cKOs mouse strains. The average area of implantation bulges was 0.077 cm^2^, 0.061 cm^2^, 0.039 cm^2^, and 0.121 cm^2^ in control, *Stat3, Gp130*, and *Lifr* cKO mice strains, respectively (Fig. 3G). The areas of the bulges in *Stat3* cKO and *Gp130* cKO were notably smaller compared to the control, whereas those in *Lifr* cKO was larger (p < 0.05). The bulges areas in *Stat3* cKO mice were markedly larger than those in *Gp130* cKO mice (Fig. 3G; p < 0.05).

Histological analysis revealed that uterine sections of all cKOs mouse exhibited the decidualized stromal cells (Fig. 4C-H). In both the control and *Lifr* cKO uterus, blastocysts were located at the antimesometrial side of the crypt (Fig. 4A, B, G, H). No blastocyst was observed in the uterus of *Stat3* and *Gp130* cKO mice (Fig. 4C, D, E, F). In the *Stat3* cKO uterus, LE was broken and shed, and the development of blood vessels was observed at implantation sites (Fig. 4D). Histological analysis of *Gp130* cKO mice uterus showed that the leukocyte infiltration and embryo death had occurred (Fig. 4F).

**Fig. 4.**
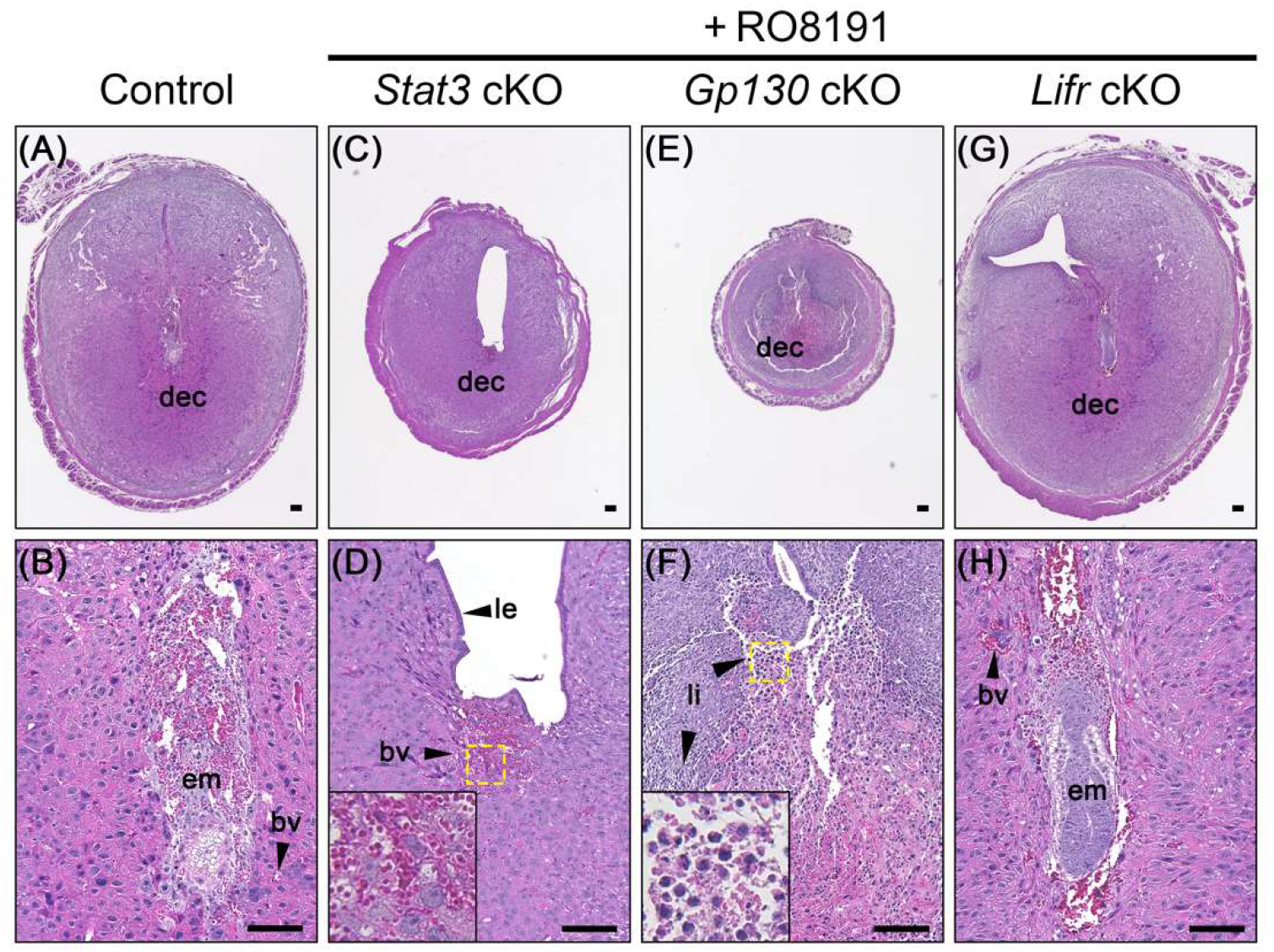
Histological characterization of uterine cross-sections from presumed implantation sites on D7. **(A, B)** Control (*Lifr*^*f/f*^) uterus after sesame oil treatment. **(C, D)** *Stat3* cKO, **(E, F)** *Gp130* cKO, and **(G, H)** *Lifr* cKO uterus after RO8191 treatment. Higher magnification in yellow frames inserted in D and F. Scale bar, 100 µm. dec: decidua; em: embryo; bv: blood vessels; le: luminal epithelium; li: leukocyte infiltration.

## Discussion

It is widely accepted that a transient elevation of E2 in the uterus can induce LIF secretion from the glandular epithelium at D4 of pregnancy in mice. LIF activates the JAK/STAT3 signaling pathway through LIFR/Gp130 and phosphorylation of STAT3 in the uterine epithelium, which is critical for embryo implantation (Cheng et al. 2001; Stewart et al. 1992). Therefore, uterine epithelial cKO mice lacking *Stat3, Gp130*, or *Lifr* exhibit defects in STAT3 signaling, resulting in implantation failure (Fukui *et al*. 2021; Hiraoka *et al*. 2020; Namiki *et al*. 2023). Although the detailed mechanisms of downstream STAT3 signaling remain elusive, rescuing implantation failure through STAT3 activation offers considerable potential for future applications.

The purpose of this study was to test whether activation of STAT3 signaling by RO8191 can initiate embryo implantation. RO8191 functions similarly to type I interferons (IFNs) as a ligand: it binds to the IFN-α receptor 2 (IFNAR2) as a homodimer, phosphorylates and activates the STAT protein family, including STAT1, STAT2, STAT3, STAT5, and STAT6, and induces IFN-inducible gene expression (Konishi et al. 2012), while type I IFNs phosphorylate STAT proteins by inducing heterodimerization of IFNAR1 and IFNAR2. In the present study, DI mice injected with RO8191 showed high implantation rates, a substantial number of implantation sites, succeed embryo development and decidualization (Fig. 1), indicating its potential as a STAT3 activator. On the other hand, individual physiological differences among the DI mice may have led to abnormal embryo development and decidualization at some implantation sites. RO8191 may bind the IFNAR2, which is known to be expressed in the luminal epithelium (Jang et al. 2017), and/or an unknown receptor to regulate gene expression, and eventually rescue embryo implantation in DI mice.

P-STAT3 was found in the uterine LE of DI mice 24 hours after RO8191 treatment (Fig. 2), while it was absent in the control, suggesting that RO8191 can directly activate the STAT3 signaling pathway in LE cells. Previous research has shown that mouse LE cells treated with LIF *in vitro* exhibited a clear single band of p-STAT3 by Western blot analysis (Cheng et al. 2001) and DI mice injected with LIF also showed a high expression level of p-STAT3 (Kobayashi et al. 2014). The p-STAT3 signal induced by RO8191 was lower compared to those induced by LIF in the previous study. It is possible that the low phosphorylation level of STAT3 in LE may be sufficient for embryo implantation and/or RO8191 activates alternative signaling pathways that controls embryo implantation independently of the STAT3 pathway.

In the present study, the uterine epithelium cKOs (*Stat3, Gp130*, and *Lifr*) also exhibited a high implantation rate and a substantial number of presumed implantation sites after RO8191 treatment (Fig. 3). This indicated that RO8191 can affect the interaction between the embryos and the uterus even in the absence of STAT3, Gp130, and LIFR in the uterine epithelium. The size of the implantation sites induced by the RO8191 administration differed among the cKOs mouse strains (Fig. 3G). The degree of the decidual reaction appears to be different, suggesting that epithelial STAT3, Gp130, and LIFR have different contribution to the decidualization. On the other hand, only *Lifr* cKO mice showed a larger size of the implantation sites compared to the control. The absence of LIFR might enhance the RO8191-induced signaling transduction, regulating the proliferation of endometrial stromal cell.

Activation of the LIF-STAT3-Egr1 signaling pathway in the endometrial stroma stimulates the decidualization of endometrial stromal cells (Liang et al. 2014). As conditional deletion of *Stat3, Gp130*, or *Lifr* in uterine epithelial cells results in defective decidualization (Catalano et al. 2005; Namiki et al. 2023; Pawar et al. 2013), LIFR/Gp130-JAK/STAT3 signaling in the uterine epithelium is indispensable for decidualization in normal pregnancy. Importantly, the present study showed that all cKOs uteri exhibited decidualized stromal cells after RO8191 treatment according to histological observations (Fig. 4). These results indicate that the activation of the STAT3 pathway by RO8191 in LE cells should play a minor role in the decidualization of uterine stromal cells. Alternatively, RO8191 should directly activate the STAT3 pathway in uterine stromal cells to initiate decidualization.

Embryo development in the decidua of *Stat3* and *Gp130* cKO uterus could not be rescued by RO8191, as no embryos were observed (Fig. 4C-F), although RO8191 exhibited low toxicity in cell culture (Konishi et al. 2012) and positive effects in *Lifr* cKO mice (Fig. 4G,H). The *Stat3* cKO uterus exhibited key features of a receptive endometrium, including partial uterine epithelial breakdown, shedding, and blood vessel development, which are essential processes for embryonic growth (Fig. 4C, D) (Whitby, Zhou, and Dimitriadis 2020). In contrast, the *Gp130* cKO did not show these characteristics. This suggested that embryonic development was already initiated but terminated at different stages of the decidualized reaction in the *Stat3* and *Gp130* cKO uterus. In the *Gp130* cKO uterus, an excessive inflammatory response and embryo death were observed after RO8191 treatment (Fig. 4F). Decidualization is typically accompanied by leukocyte infiltration, with approximately 15% of uterine decidual cells being immune cells, primarily uterine NK cells, macrophages, and T lymphocytes (Erlebacher 2013). However, abnormal leukocyte infiltration also can lead to the early embryo loss (Baines et al. 1997). Previous research has shown that the absence of epithelial Gp130 reduces the expression of the progesterone receptor and ALOX15, a downstream target of the progesterone receptor (Namiki et al. 2023). This reduction leads to defective decidualization and an excessive inflammatory response in *Gp130* cKO mice at D4 (Namiki et al. 2023). This indicates that RO8191 was ineffective in alleviating the defective decidualization and excessive inflammation in *Gp130* cKO mice. The marked failure of decidualization in the *Stat3* and *Gp130* cKO uteri suggests that STAT3 and Gp130 play crucial but distinct roles in the RO8191-mediated signaling, which regulates decidualization and embryonic development.

RO8191 also has the advantage of convenient oral administration in mice, which is patient-friendly for the future application (Konishi et al. 2012). In this study, we demonstrated that RO8191 can contribute to embryo implantation in mice. Although the exact mechanism of RO8191 on implantation remains unclear, it is promising that RO8191 has potential for the treatment of implantation failure.

## Acknowledgements

We thank Dr. T. Daikoku (Kanazawa University) and Dr. S.K. Dey (Cincinnati Children’s Hospital Medical Center) for providing *Ltf*^*iCre*^ mice, Dr. S. Akira (Osaka University) for providing *Stat3*^*flox*^ mice, and Dr. W. Muller (University of Manchester) for providing *Gp130*^*flox*^ mice. We also thank Medical Research Council (MRC) for providing *Lifr*^*tm1a(EUCOMM)Hmgu*^. FLPe transgenic mouse strain (RBRC01834) was provided by RIKEN BRC through the National BioResource Project of the MEXT/AMED, Japan. This work was supported by Grants-in-Aid for Scientific Research from the JSPS (23K23785 to E.H.). This work was partially supported by the Center for Human and Animal Symbiosis Science, Azabu University, and a research project grant awarded by the Azabu University Research Services Division.

## References

Baines, M. G., A. J. Duclos, E. Antecka, and E. K. Haddad. 1997. “Decidual Infiltration and Activation of Macrophages Leads to Early Embryo Loss.” American Journal of Reproductive Immunology (New York, N.Y.: 1989) 37(6):471–77. doi: 10.1111/j.1600-0897.1997.tb00262.x.

Betz, Ulrich A. K., Wilhelm Bloch, Maries van den Broek, Kanji Yoshida, Tetsuya Taga, Tadamitsu Kishimoto, Klaus Addicks, Klaus Rajewsky, and Werner Müller. 1998. “Postnatally Induced Inactivation of Gp130 in Mice Results in Neurological, Cardiac, Hematopoietic, Immunological, Hepatic, and Pulmonary Defects.” The Journal of Experimental Medicine 188(10):1955–65.

Catalano, Rob D., Martin H. Johnson, Elizabeth A. Campbell, D. Stephen Charnock-Jones, Stephen K. Smith, and Andrew M. Sharkey. 2005. “Inhibition of Stat3 Activation in the Endometrium Prevents Implantation: A Nonsteroidal Approach to Contraception.” Proceedings of the National Academy of Sciences of the United States of America 102(24):8585–90. doi: 10.1073/pnas.0502343102.

Cheng, Jr-Gang, Jim Ray Chen, Lidia Hernandez, W. Greg Alvord, and Colin L. Stewart. 2001. “Dual Control of LIF Expression and LIF Receptor Function Regulate Stat3 Activation at the Onset of Uterine Receptivity and Embryo Implantation.” Proceedings of the National Academy of Sciences 98(15):8680– 85. doi: 10.1073/pnas.151180898.

Cheng, JrGang, Gracy Rosario, Tatiana V. Cohen, Jianbo Hu, and Colin L. Stewart. 2017. “Tissue-Specific Ablation of the LIF Receptor in the Murine Uterine Epithelium Results in Implantation Failure.” Endocrinology 158(6):1916–28. doi: 10.1210/en.2017-00103.

Coughlan, C., W. Ledger, Q. Wang, Fenghua Liu, Aygul Demirol, Timur Gurgan, R. Cutting, K. Ong, H. Sallam, and T. C. Li. 2014. “Recurrent Implantation Failure: Definition and Management.” Reproductive BioMedicine Online 28(1):14–38. doi: 10.1016/j.rbmo.2013.08.011.

Daikoku, Takiko, Yuya Ogawa, Jumpei Terakawa, Akiyo Ogawa, Tony DeFalco, and Sudhansu K. Dey. 2014. “Lactoferrin-iCre: A New Mouse Line to Study Uterine Epithelial Gene Function.” Endocrinology 155(7):2718–24. doi: 10.1210/en.2014-1265.

Erlebacher, Adrian. 2013. “Immunology of the Maternal-Fetal Interface.” Annual Review of Immunology 31(Volume 31, 2013.:387–411. doi: 10.1146/annurev-immunol-032712-100003.

Fukui, Yamato, Yasushi Hirota, Tomoko Saito-Fujita, Shizu Aikawa, Takehiro Hiraoka, Tetsuaki Kaku, Tomoyuki Hirata, Shun Akaeda, Mitsunori Matsuo, Ryoko Shimizu-Hirota, Norihiko Takeda, Masahito Ikawa, and Yutaka Osuga. 2021. “Uterine Epithelial LIF Receptors Contribute to Implantation Chamber Formation in Blastocyst Attachment.” Endocrinology 162(11):bqab169. doi: 10.1210/endocr/bqab169.

Giudice, L. C. 1999. “Potential Biochemical Markers of Uterine Receptivity.” Human Reproduction (Oxford, England) 14 Suppl 2:3–16. doi: 10.1093/humrep/14.suppl_2.3.

Hiraoka, Takehiro, Yasushi Hirota, Yamato Fukui, Mona Gebril, Tetsuaki Kaku, Shizu Aikawa, Tomoyuki Hirata, Shun Akaeda, Mitsunori Matsuo, Hirofumi Haraguchi, Mayuko Saito-Kanatani, Ryoko Shimizu-Hirota, Norihiko Takeda, Osamu Yoshino, Tomoyuki Fujii, and Yutaka Osuga. 2020. “Differential Roles of Uterine Epithelial and Stromal STAT3 Coordinate Uterine Receptivity and Embryo Attachment.” Scientific Reports 10(1):15523. doi: 10.1038/s41598-020-72640-0.

Hu, Wenwei, Zhaohui Feng, Angelika K. Teresky, and Arnold J. Levine. 2007. “P53 Regulates Maternal Reproduction through LIF.” Nature 450(7170):721–24. doi: 10.1038/nature05993.

Jang, Hwanhee, Yohan Choi, Inkyu Yoo, Jisoo Han, Minjeong Kim, and Hakhyun Ka. 2017. “Characterization of Interferon α and β Receptor IFNAR1 and IFNAR2 Expression and Regulation in the Uterine Endometrium during the Estrous Cycle and Pregnancy in Pigs.” Theriogenology 88:166–73. doi: 10.1016/j.theriogenology.2016.09.025.

Kanki, Hiroaki, Hitomi Suzuki, and Shigeyoshi Itohara. 2006. “High-Efficiency CAG-FLPe Deleter Mice in C57BL/6J Background.” Experimental Animals 55(2):137–41. doi: 10.1538/expanim.55.137.

Kliman, Harvey J., and David Frankfurter. 2019. “Clinical Approach to Recurrent Implantation Failure: Evidence-Based Evaluation of the Endometrium.” Fertility and Sterility 111(4):618–28. doi: 10.1016/j.fertnstert.2019.02.011.

Kobayashi, Ryosuke, Jumpei Terakawa, Yasumasa Kato, Shafiqullah Azimi, Naoko Inoue, Yasushige Ohmori, and Eiichi Hondo. 2014. “The Contribution of Leukemia Inhibitory Factor (LIF) for Embryo Implantation Differs among Strains of Mice.” Immunobiology 219(7):512–21. doi: 10.1016/j.imbio.2014.03.011.

Kojima, Yoji, Oliver H. Tam, and Patrick P. L. Tam. 2014. “Timing of Developmental Events in the Early Mouse Embryo.” Seminars in Cell & Developmental Biology 34:65–75. doi: 10.1016/j.semcdb.2014.06.010.

Konishi, Hideyuki, Koichi Okamoto, Yusuke Ohmori, Hitoshi Yoshino, Hiroshi Ohmori, Motooki Ashihara, Yuichi Hirata, Atsunori Ohta, Hiroshi Sakamoto, Natsuko Hada, Asao Katsume, Michinori Kohara, Kazumi Morikawa, Takuo Tsukuda, Nobuo Shimma, Graham R. Foster, William Alazawi, Yuko Aoki, Mikio Arisawa, and Masayuki Sudoh. 2012. “An Orally Available, Small-Molecule Interferon Inhibits Viral Replication.” Scientific Reports 2:259. doi: 10.1038/srep00259.

Liang, Xiao-Huan, Wen-Bo Deng, Ming Li, Zhen-Ao Zhao, Tong-Song Wang, Xu-Hui Feng, Yu-Jing Cao, En-Kui Duan, and Zeng-Ming Yang. 2014. “Egr1 Protein Acts Downstream of Estrogen-Leukemia Inhibitory Factor (LIF)-STAT3 Pathway and Plays a Role during Implantation through Targeting Wnt4 *.” Journal of Biological Chemistry 289(34):23534–45. doi: 10.1074/jbc.M114.588897.

Macklon, N. S., J. P. M. Geraedts, and B. C. J. M. Fauser. 2002. “Conception to Ongoing Pregnancy: The ‘Black Box’ of Early Pregnancy Loss.” Human Reproduction Update 8(4):333–43. doi: 10.1093/humupd/8.4.333.

Namiki, Takafumi, Jumpei Terakawa, Harumi Karakama, Michiko Noguchi, Hironobu Murakami, Yoshinori Hasegawa, Osamu Ohara, Takiko Daikoku, Junya Ito, and Naomi Kashiwazaki. 2023. “Uterine Epithelial Gp130 Orchestrates Hormone Response and Epithelial Remodeling for Successful Embryo Attachment in Mice.” Scientific Reports 13(1):854. doi: 10.1038/s41598-023-27859-y.

Onishi, Kento, and Peter W. Zandstra. 2015. “LIF Signaling in Stem Cells and Development.” Development 142(13):2230–36. doi: 10.1242/dev.117598.

Pastuschek, Jana, Jenny Poetzsch, Diana M. Morales-Prieto, Ekkehard Schleußner, Udo R. Markert, and Georgi Georgiev. 2015. “Stimulation of the JAK/STAT Pathway by LIF and OSM in the Human Granulosa Cell Line COV434.” Journal of Reproductive Immunology 108:48–55. doi: 10.1016/j.jri.2015.03.002.

Pawar, Sandeep, Elina Starosvetsky, Grant D. Orvis, Richard R. Behringer, Indrani C. Bagchi, and Milan K. Bagchi. 2013. “STAT3 Regulates Uterine Epithelial Remodeling and Epithelial-Stromal Crosstalk During Implantation.” Molecular Endocrinology 27(12):1996–2012. doi: 10.1210/me.2013-1206.

Robb, Lorraine, Eva Dimitriadis, Ruili Li, and Lois A. Salamonsen. 2002. “Leukemia Inhibitory Factor and Interleukin-11: Cytokines with Key Roles in Implantation.” Journal of Reproductive Immunology 57(1):129–41. doi: 10.1016/S0165-0378(02)00012-8.

Salleh, Naguib. 2014. “Diverse Roles of Prostaglandins in Blastocyst Implantation.” The Scientific World Journal 2014:e968141. doi: 10.1155/2014/968141.

Song, Haengseok, Hyunjung Lim, Sanjoy K. Das, Bibhash C. Paria, and Sudhansu K. Dey. 2000. “Dysregulation of EGF Family of Growth Factors and COX-2 in the Uterus during the Preattachment and Attachment Reactions of the Blastocyst with the Luminal Epithelium Correlates with Implantation Failure in LIF-Deficient Mice.” Molecular Endocrinology 14(8):1147–61. doi: 10.1210/mend.14.8.0498.

Stewart, Colin L., Petr Kaspar, Lisa J. Brunet, Harshida Bhatt, Inder Gadi, Frank Köntgen, and Susan J. Abbondanzo. 1992. “Blastocyst Implantation Depends on Maternal Expression of Leukaemia Inhibitory Factor.” Nature 359(6390):76–79. doi: 10.1038/359076a0.

Takeda, K., T. Kaisho, N. Yoshida, J. Takeda, T. Kishimoto, and S. Akira. 1998. “Stat3 Activation Is Responsible for IL-6-Dependent T Cell Proliferation through Preventing Apoptosis: Generation and Characterization of T Cell-Specific Stat3-Deficient Mice.” Journal of Immunology (Baltimore, Md.: 1950) 161(9):4652– 60.

Terakawa, Jumpei, Shoichi Wakitani, Makoto Sugiyama, Naoko Inoue, Yasushige Ohmori, Yasuo Kiso, Yoshinao Z. Hosaka, and Eiichi Hondo. 2011. “Embryo Implantation Is Blocked by Intraperitoneal Injection with Anti-LIF Antibody in Mice.” Journal of Reproduction and Development 57(6):700–707. doi: 10.1262/jrd.11-048H.

Wang, Haibin, and Sudhansu K. Dey. 2006. “Roadmap to Embryo Implantation: Clues from Mouse Models.” Nature Reviews Genetics 7(3):185–99. doi: 10.1038/nrg1808.

White, Christine A., Jian-Guo Zhang, Lois A. Salamonsen, Manuel Baca, W. Douglas Fairlie, Donald Metcalf, Nicos A. Nicola, Lorraine Robb, and Evdokia Dimitriadis. 2007. “Blocking LIF Action in the Uterus by Using a PEGylated Antagonist Prevents Implantation: A Nonhormonal Contraceptive Strategy.” Proceedings of the National Academy of Sciences of the United States of America 104(49):19357–62. doi: 10.1073/pnas.0710110104.

Yang, Z. M., S. P. Le, D. B. Chen, J. Cota, V. Siero, K. Yasukawa, and M. J. Harper. 1995. “Leukemia Inhibitory Factor, LIF Receptor, and Gp130 in the Mouse Uterus during Early Pregnancy.” Molecular Reproduction and Development 42(4):407–14. doi: 10.1002/mrd.1080420406.

